# A viral-specific CD4^+^ T cell response protects female mice from Coxsackievirus B3 infection

**DOI:** 10.1101/2023.10.24.563774

**Authors:** Aryamav Pattnaik, Adeeba H. Dhalech, Stephanie A. Condotta, Caleb Corn, Martin J. Richer, Laura M. Snell, Christopher M. Robinson

## Abstract

Biological sex plays an integral role in the immune response to various pathogens. The underlying basis for these sex differences is still not well defined. Here, we show that Coxsackievirus B3 (CVB3) induces a viral-specific CD4^+^ T cell response that can protect female mice from mortality. We found that CVB3 can induce expansion of CD62L^lo^ CD4^+^ T cells in the mesenteric lymph node and spleen of female but not male mice as early as 5 days post-inoculation, indicative of activation. Using a recombinant CVB3 virus expressing a model CD4^+^ T cell epitope, we found that this response is due to viral antigen and not bystander activation. Finally, the depletion of CD4^+^ T cells before infection increased mortality in female mice, indicating that CD4^+^ T cells play a protective role against CVB3 in our model. Overall, these data demonstrated that CVB3 can induce an early CD4 response in female but not male mice and further emphasize how sex differences in immune responses to pathogens affect disease outcomes.

## 1. Introduction

Biological sex is critical in the immune response to various pathogens (1, 2). Generally, women exhibit a more robust immune response to infections than men, and women often develop higher antibody responses to vaccines (3). This sex bias is likely due to differences in the number and activation of immune cells. For example, women have higher CD4^+^ to CD8^+^ T cell ratios than men (1). Also, T cells from females have enriched expression of genes involved in T cell activation (4). Furthermore, women exhibit stronger type I interferon responses, and female antigen-presenting cells are more efficient in presenting peptides than their male counterparts (5-7). Therefore, females may be inherently primed to mount more robust immune responses that limit pathogen replication and pathogenesis better than males. However, this comes at a cost, as females frequently experience more severe infection symptoms, more adverse vaccine reactions, and are more vulnerable to autoimmune disorders (2). Unfortunately, the mechanism dictating the sex bias in immunity remains elusive. Thus, knowing how sex impacts immune responses to infections and vaccines is crucial for therapeutic development and to aid in vaccine strategies to target both men and women.

Enteroviruses continue to pose a substantial global public health issue (8-11). These viruses are a group of positive-sense stranded RNA viruses classified in the Picornavirus family. Among enteroviruses, Coxsackievirus B3 (CVB3) is commonly isolated during surveillance and causes viral myocarditis and aseptic meningitis (12, 13). CVB3 is transmitted through the fecal-oral route and initiates infection in the intestine. Sex is known to be a pivotal contributor to CVB3 infections in humans. Men are two to three times more likely to develop myocarditis than women (14). This disparity is also mimicked in mouse models of CVB3 infection. Immune differences between males and females have been implicated in the pathology of CVB3-induced myocarditis; however, the mechanisms leading to these events are still largely unclear.

Data from C57BL/6 mouse models show that acute CVB3-induced myocarditis is characterized by inflammatory infiltration of immune cells, including CD4^+^ and CD8^+^ T cells (15, 16). Here, T cells likely contribute to viral clearance; however, other mouse models indicate that T cells can also contribute to CVB3-induced disease (17-20). In infection of Balb/c mice, CVB3 induces autoimmune-mediated myocarditis. In this model, differences in the CD4^+^ T helper (Th) 1 and Th2 response following infection contribute to the sex bias in myocarditis. Following CVB3 infection, males mount a predominantly Th1, which enhances acute inflammation and myocarditis (18, 20). In contrast, females skew towards a Th2 response that can limit inflammation in the heart. The mechanism for this difference is still unclear, but sex hormones can contribute to the Th1/Th2 balance following infection (21). Moreover, γδ T cells and CD4^+^ T regulatory cells (Tregs) can also promote or limit disease (22-25). Overall, these data indicate that T cells largely contribute to disease, yet how CD4^+^ T cells are activated in acute CVB3 infection and how these T cells contribute to cell-mediated viral clearance is incompletely understood.

Research from our lab and many others have shown that sex hormones influence the pathogenesis of CVB3 in mice (21, 26-29). Using an oral inoculation mouse model, our lab has demonstrated that gonadectomy impacts intestinal CVB3 replication and dissemination in male and female mice (26, 27). Castration of male mice also completely protected males from mortality following CVB3 infection, and exogenous testosterone treatment restored lethality. This suggests that testosterone plays a vital role in the mortality of male mice infected with CVB3. However, the impact of testosterone is complex since exogenous testosterone treatment in female mice did not increase mortality. These data indicate other immune cells may protect female mice from CVB3 infection. In support of this, we recently identified a sex difference in the T cell response to CVB3. We demonstrated that activated, viral-specific CD8^+^ T cells in female but not male mice expand and protect against CVB3 infection (30). Similarly, we found that CD4^+^ T cells expand in a sex-dependent manner. Here, we sought to explore further the CD4^+^ T cell response to CVB3 infection in male and female mice. Our data indicate that CD4^+^ T cells are activated in female but not male mice as early as 5 post-inoculation (dpi), and viral-specific CD4^+^ T cells expand in females but not males. Finally, we show that CD4^+^ T cells are protective against CVB3 infections in female mice. These data add to the growing body of evidence highlighting the existence of sex-specific immune regulation in response to viral infections.

## 2. Materials and Methods

### 2.1. Cells and virus

HeLa cells were cultivated in Dulbecco’s modified Eagle’s medium (DMEM) supplemented with 10% calf serum and 1% penicillin-streptomycin. The cells were maintained at 37°C in an environment containing 5% CO2. The infectious clones CVB3-Nancy and recombinant CVB3-H3 were obtained from Marco Vignuzzi at the Pasteur Institute in Paris, France. and transfected in HeLa cells as previously described (26). A standard plaque assay with HeLa cells was used to quantify the virus. The rCVB3.6 was graciously provided to us by Lindsay Whitton and Taishi Kimura (Scripps Research Institute in La Jolla, California) and passaged in HeLa cells.

### 2.2. Mouse experiments

C57BL/6, *PVR^+/+^, Ifnar-/-* mice were obtained from S. Koike in Tokyo, Japan (31, 32). Age-matched mice, ranging in age from 8 to 14 weeks at the time of infection, were used for all experiments. Male and female mice were orally inoculated with 5x10^7^ PFUs or intraperitoneally inoculated with 1x10^4^ PFUs of CVB3-Nancy. For rCVB3.6 experiments, mice were intraperitoneally inoculated with 1x10^4^ PFU of CVB3-H3 as control or 5x10^7^-1x10^8^ PFU of rCVB3.6. Data from the mouse investigations were combined from two to three experiments, with each group consisting of at least three mice per study.

### 2.3. SMARTA T cells and adoptive transfer

Transgenic T cells with TCRs specific to LCMV GP_61-80_ peptide were isolated from SMARTA mice, which have been previously characterized and were bred on a CD45.1 background (33). SMARTA T cells (CD45.1) were adoptively transferred into our *Ifnar^-/-^*(CD45.2) mouse model i.v. via the retro-orbital sinus one day before infection. Female *Ifnar^-/-^* mice were then ip inoculated with 10^8^ PFU of rCVB3.6 or 10^4^ PFU of wild-type CVB3 (wtCVB3) as a control. As a control for gating GP_66_-specific CD4 T cells, we also infected female *Ifnar^-/-^*mice with LCMV (data not shown). The spleen was harvested from infected mice at 5dpi, and virus-specific SMARTA CD4 T cells were analyzed by flow cytometry by expression of CD45.1.

### 2.4. Flow cytometry analysis

At indicated time points post-infection, the spleen and mesenteric lymph nodes from male and female mice were collected and mechanically disrupted to obtain a single-cell suspension. RBC lysis buffer from BioLegend (catalog # 420301) was used to remove erythrocytes. Afterward, the cells were washed and incubated with TruStain fcX (CD16/CD32, Clone 93, BioLegend, catalog # 101320) to prevent non-specific binding, and the immune cells of interest were then stained with appropriate surface antibodies. The samples were subjected to analysis using a BD LSRFortessa flow cytometer in combination with FlowJo software (BD Biosciences). The following mouse antibodies, in various fluorochrome combinations, were utilized for the staining: CD4 (clone GK1.5, BioLegend, catalog #100412, #100406), CD8a (clone 53-6.7, BioLegend, catalog, #100707 #100711), CD11a (clone M17/4, BioLegend, catalog # 101106), CD62L (clone MEL-14, BioLegend, catalog #104438), CD49d (clone R1-2, BioLegend, catalog #103618).

### 2.5. CD4^+^ T cell depletion

Two days before infection, CD4^+^ T cells were depleted in male and female mice by intraperitoneally injecting mice with 200 μg of an anti-CD4 antibody (anti-mouse CD4 clone: GK1.5, BioXcell, catalog # BE0003-1). A group of mice receiving an intraperitoneal injection of an isotype control antibody (rat IgG2b LTF-2, BioXcell, catalog # BE0090) was used as a control. The following day, a tail vein blood sample was collected from each mouse. The blood samples were then stained and analyzed by flow cytometry to confirm the successful depletion of CD4^+^ T cells.

### 2.6. Statistical Analysis

Comparisons between the control and study groups were analyzed using either an unpaired t-test or a one-way analysis of variance (ANOVA), depending on the experimental design. To visualize the variability, error bars in the figures were plotted to represent the standard errors of the means. A significance level of p < 0.05 was used to determine if there were any meaningful differences between the groups. All statistical analyses and graphs were generated using GraphPad Prism 10 (GraphPad Software, La Jolla, CA), ensuring accurate and comprehensive data representation.

## 3. Results

### 3.1. Antigen-experienced CD4^+^ T cells expand in the mesenteric lymph nodes of CVB3-infected female but not male mice

We previously found that following oral inoculation, CVB3 induces a sex-dependent expansion of T cells in the spleen (30). However, since CVB3 initiates infection in the intestine, we examined the CD4^+^ T cell response in the local lymph node following infection. Male and female *Ifnar^-/-^*mice were orally inoculated with 5x10^7^ PFUs of CVB3, and the mesenteric lymph nodes (MLNs) were harvested at 5dpi. We chose to examine immune cells at 5dpi because we have previously shown that mortality in male mice typically begins at this time point (26, 27). Following CVB3 infection, we found that the number of CD4^+^ T cells showed a trending increase in the infected female mice compared to uninfected female mice (Fig. 1A and Fig. 1B). However, this increase was not statistically significant (p=0.0816). In contrast, we observed a significant increase in the number of CD4^+^ T cells in infected female *Ifnar^-/-^* mice compared to infected male mice (Fig. 1B). Next, we assessed the CD4^+^ T cell activation by the expression of CD62L. Naïve T cells are CD62L^hi^ and activated T cells differentiate into effector subtypes during acute infections. During this effector phase, CD62L is downregulated (34-36). Following oral inoculation, we found a significant increase in the number of CD62L^lo^ CD4^+^ T cells in infected female mice compared to uninfected female mice and infected male mice (Fig. 1C). Moreover, this increase was sex-dependent and not observed between infected and uninfected male mice.

**Figure 1.**
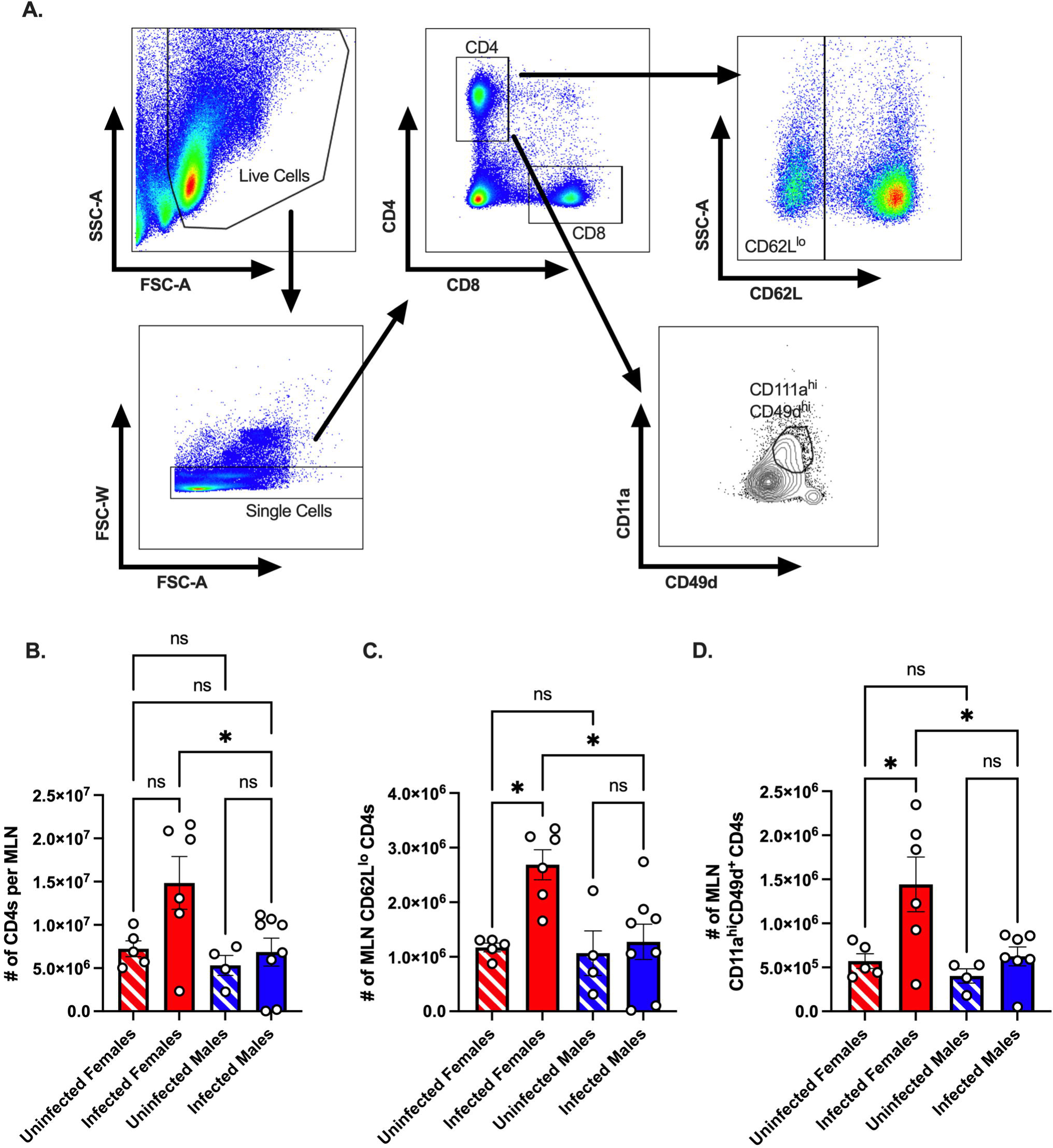
The CD4^+^ T cell response in the mesenteric lymph node of CVB3-infected *Ifnar^-/-^*mice. Male and female *Ifnar^-/-^* mice were orally inoculated with 5x10^7^ PFUs of CVB3. The mesenteric lymph nodes (MLN) were harvested at 5dpi and processed for analysis by flow cytometry. (A) Representative gating strategy for CD4^+^ T cells, CD62L^lo^ CD4^+^ T cells, and CD11a^hi^CD49d^+^ CD4 T cells. (B) The number of MLN CD4^+^ T cells in male and female mice 5dpi. (C) The number of CD62L^lo^ CD4^+^ T cells in the MLNs at 5dpi. (D) The number of MLN CD11a^hi^CD49d^+^ CD4 T cells. Data points represent individual mice. Data are from at least two independent experiments. *p<0.05, One-way ANOVA.

Next, the downregulation of CD62L on CD4^+^ T cells can occur through bystander activation rather than direct antigen-specific engagement of the T cell receptor (37). Previous studies have established that antigen-experienced T cells can be followed regardless of their specificity using surrogate markers (34, 38-42). To investigate if CVB3-induced virus-specific CD4^+^ T cells in the mesenteric lymph nodes, we measured the expression of CD11a and CD49d on CD4^+^ T cells as a measure of antigen-experienced CD4^+^ T cells (Fig. 1A). We found a significant increase in the number of CD11a^hi^CD49d^+^ CD4^+^ T cells in infected females compared to uninfected females (Fig. 1D). In contrast, no difference in the number of CD11a^hi^CD49d^+^ CD4^+^ T cells between uninfected and infected males was observed. Moreover, the number of CD11a^hi^CD49d^+^ CD4^+^ T cells in infected female mice was significantly higher than in infected male mice. Taken together, these data indicate that CVB3 induces activated, antigen-experienced CD4^+^ cells at 5dpi in mesenteric lymph nodes of female but not male *Ifnar^-/-^* mice.

### 3.2. Sex-dependent activation of CD4^+^ T cells in the spleen occurs as early as 5 dpi in CVB3-infected mice

We previously found that splenic CD4^+^ and CD8^+^ T cells expand at 5dpi in orally CVB3-infected female *Ifnar^-/-^* mice but not infected male *Ifnar^-/-^* mice (30). In contrast to antigen-experienced CD8^+^ T cells that increase as early as 5 dpi, we found that splenic antigen-experienced CD4^+^ T cells from female mice do not increase until 15 dpi (30). However, we hypothesized that splenic CD4^+^ T cells may be undergoing signs of early activation even though the upregulation of the cell surface expression of the surrogate makers, CD11a and CD49d, had not occurred. Therefore, to test this hypothesis, we assessed CD4^+^ T cell activation in the spleen by the expression of CD62L. Male and female *Ifnar^-/-^* mice were orally inoculated with 5x10^7^ PFUs of CVB3, and the spleen from infected and uninfected mice was harvested at 5 dpi. Following CVB3 inoculation, we observed a significant increase in the number of CD62L^lo^ CD4^+^ T cells in infected female mice compared to infected male mice at 5dpi (Fig. 2A). Further, infected female mice had significantly higher numbers of CD62L^lo^ CD4^+^ T cells than uninfected female mice. In contrast, no difference was observed in the number of CD62L^lo^ CD4^+^ T cells between uninfected and infected male mice. Next, to confirm that the CD4^+^ T cells were becoming activated in female mice but not male mice, we examined if the CD4^+^ T cells were undergoing blast transformation. Within a few hours of antigen activation, CD4^+^ T cells can undergo blast transformation, which results in increased granularity as they develop into mature effector cells (43, 44). To measure changes in granularity, we assessed the mean fluorescence intensity (gMFI) for the side scatter area (SSC) of the CD62L^lo^ CD4^+^ T cell population. We found that CD62L^lo^ CD4^+^ T cells from infected female mice had a significant increase in granularity, as measured by the SSC, compared to CD62L^lo^ CD4^+^ T cells from uninfected female mice (Fig. 2B). Overall, these data indicate that CVB3 induces early activation of CD4^+^ T cells at 5 dpi in female mice but not male mice.

**Figure 2.**
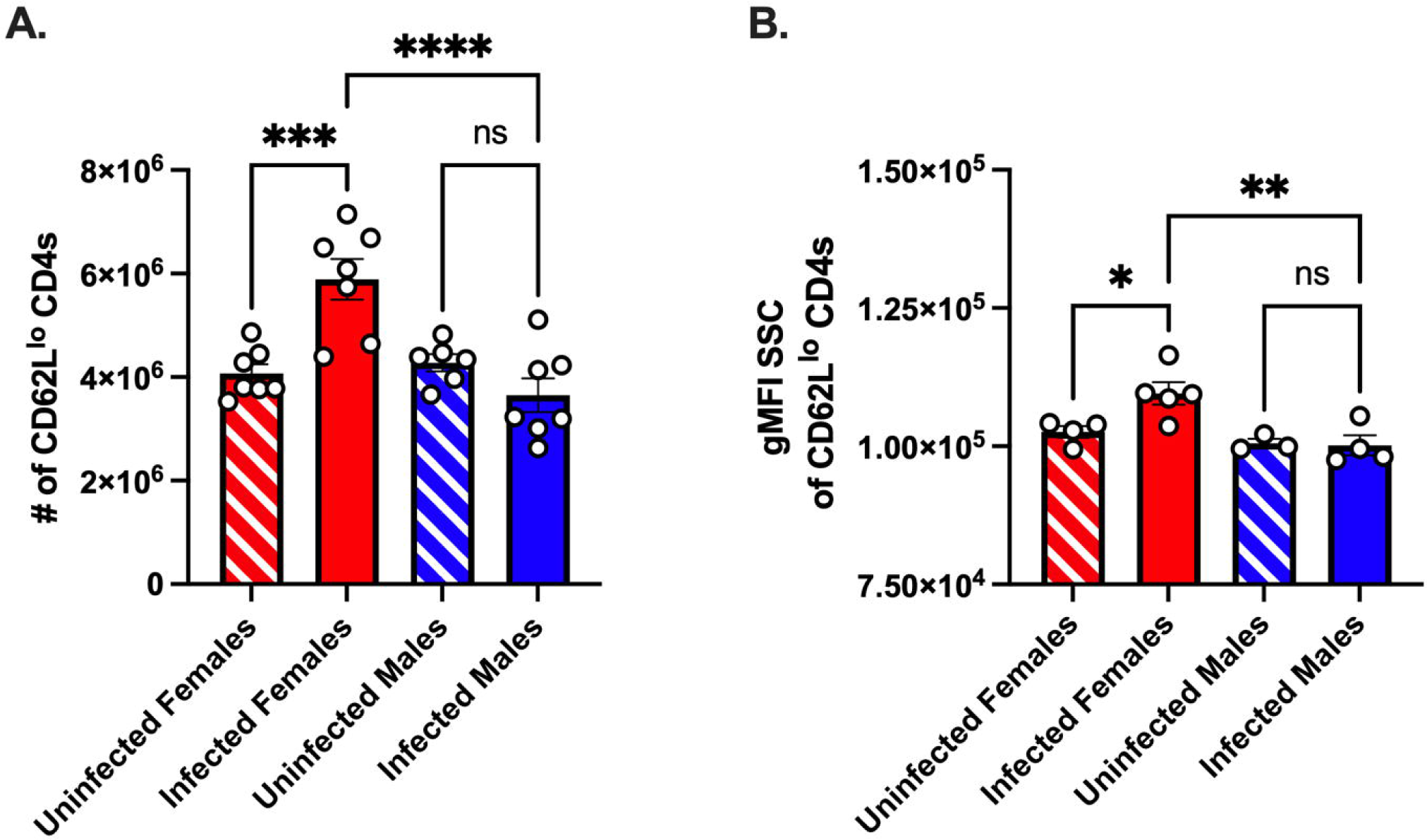
CVB3 induces early expansion of activated splenic CD4^+^ T cells in female *Ifnar^-/-^* mice. (A) The number of CD62L^lo^ CD4^+^ T cells in the spleen 5dpi in *Ifnar^-/-^*mice. (B) Representative of the gMFI SSC of CD62L^lo^ CD8^+^ T cells from two independent experiments. Data points represent individual mice. Data are from two independent experiments.*p<0.05, **p<0.01, ***p<0.001, ****p<0.0001 One-way ANOVA.

### 3.3. CVB3 induces viral-specific splenic CD4^+^ T cells as early as five days post-inoculation

Unfortunately, few studies have identified immunodominant T cell epitopes for CVB3. Therefore, to determine if the CD4^+^ T cell response was viral-specific, we used a recombinant CVB3 (rCVB3.6) that encodes a well-characterized CD4^+^ T cell epitope from lymphocytic choriomeningitis virus (LCMV) (Fig. 3A). This recombinant CVB3, while attenuated *in vivo*, still leads to productive infection, generating high tissue titers, and is cleared similarly to wild-type CVB3 (17, 45, 46). We ip inoculated female *Ifnar^-/-^* mice with rCVB3.6 or wild-type CVB3 (wtCVB3) as a control. The spleen was harvested from infected mice at 15 dpi, and virus-specific CD4^+^ T cells were analyzed by flow cytometry using an H2-D^b^ GP_66_ tetramer. We initially chose to study CD4^+^ T cells at 15 dpi, based on our previous data measuring the expansion of antigen-experienced CD4^+^ T cells (30). At 15 dpi, we observed a significant increase in the number of GP_66_-specific CD4^+^ T cells in the spleen from female mice infected with the rCVB3.6 virus compared to female mice infected with wtCVB3 (Fig. 3B). We observed a significant increase in the number of GP_66_^+^CD62L^lo^ CD4^+^ T cells from female mice inoculated with rCVB3.6 compared to wtCVB3 infected mice (Fig. 3C). Overall, these data indicate that CVB3 drives the expansion of activated, virus-specific CD4^+^ T cells at 15 dpi in female mice.

**Figure 3.**
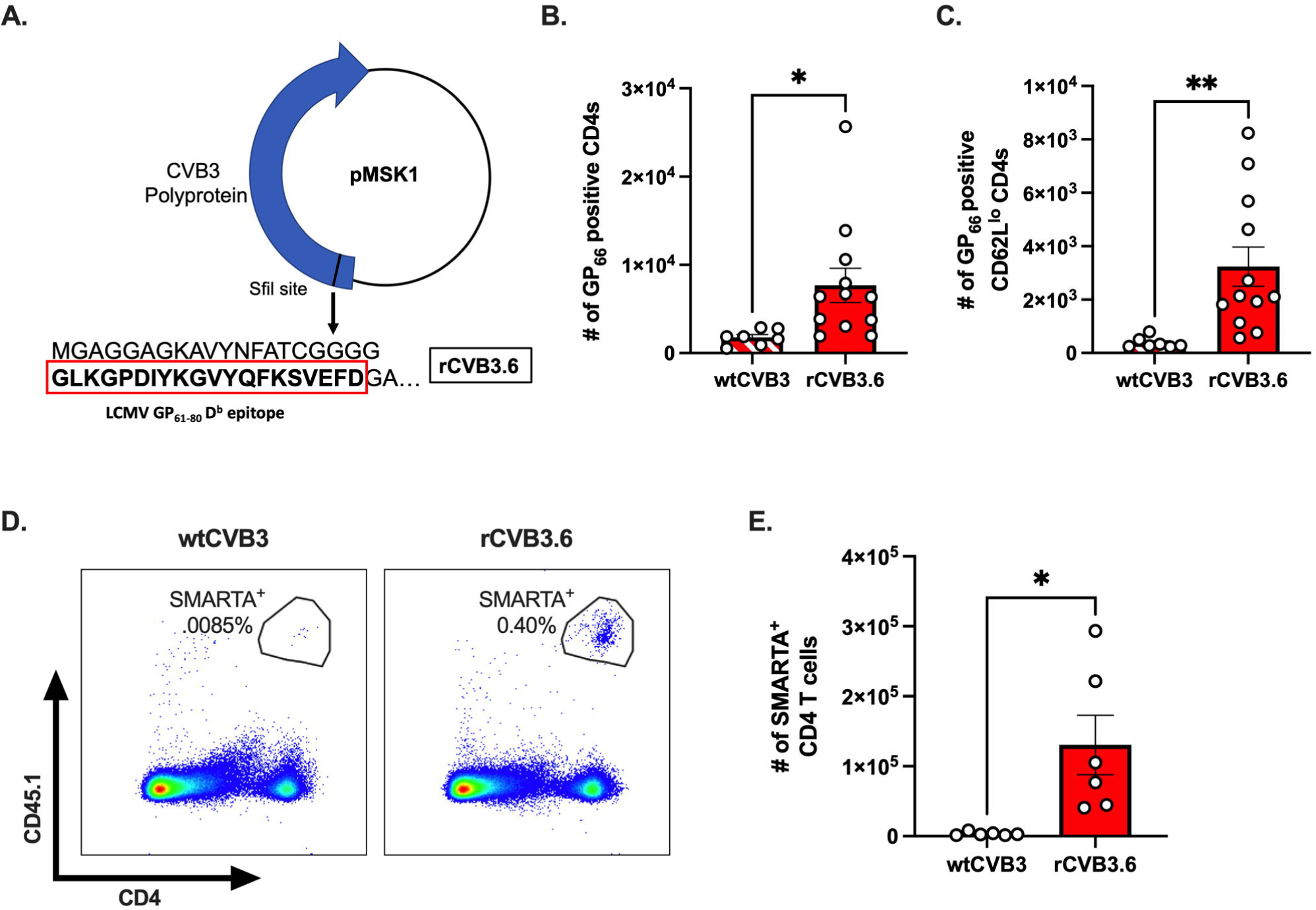
CVB3 induces a virus-specific expansion of CD4^+^ T cells in female mice. (A) The rCVB3.6 amino acid sequence encoding the LCMV GP_61-80_ CD4^+^ T cell epitope (adapted from (17)). (B) The number of splenic CD4^+^ T cells detected with H2-D^b^ GP_66_ tetramer in female *Ifnar^-/-^* mice 15dpi. (C) The number of H2-D^b^ GP_66_ tetramer positive splenic CD62L^lo^ CD4^+^ T cells. (D) Representative gating strategy for SMARTA^+^ T cells, based on CD45.1 expression, following infection with wtCVB3 or rCVB3.6. (E) The number of splenic SMARTA^+^ T cells in female mice infected with wtCVB3 or rCVB3.6. All data are from two independent experiments and are shown as mean ± SEM. *p<0.05, unpaired t-test.

Next, since we observed markers of activation on CD4^+^ T cells in the spleen as early as 5 dpi, we reasoned that the expression of surrogate markers may not detect early virus-specific CD4s. Therefore, we adoptively transferred SMARTA T cells into our mouse model to enhance our sensitivity. T cells from SMARTA mice have a transgenic T cell receptor for the LCMV GP_61-80_ T cell epitope encoded in the rCVB3.6 (47, 48). Following the adoptive transfer of SMARTA T cells, we ip inoculated mice with either rCVB3.6 or wtCVB3 and harvested the spleen 5 dpi. We found a significant increase in SMARTA CD4 T cells in the spleen of female mice inoculated with rCVB3.6 compared to wtCVB3 (Fig. 3D and 3E). These data indicate that virus-specific CD4^+^ T cells are expanding in the spleen of infected mice as early as 5 dpi in ip inoculated female mice.

### 3.4. CD4^+^ T cells protect female mice but not male mice from CVB3-induced lethality

We have previously shown that female *Ifnar^-/-^* mice are protected against CVB3-induced mortality (26, 27). Since CD4^+^ T cells expand in female mice following infection, we hypothesized that these T cells might offer protection against CVB3-induced lethality. To test this hypothesis, we depleted CD4^+^ T cells with a monoclonal antibody before CVB3 inoculation (Fig. 4A and 4B). Following CVB3 infection, we observed a significant increase in mortality in female mice depleted of CD4^+^ T cells compared to female mice treated with an isotype control (Fig. 4C). In contrast to females, depletion of CD4^+^ T cells in males did not significantly enhance mortality (Fig. 4D). These data demonstrate that CD4^+^ T cells play a significant role in protection against CVB3 in female mice.

**Figure 4.**
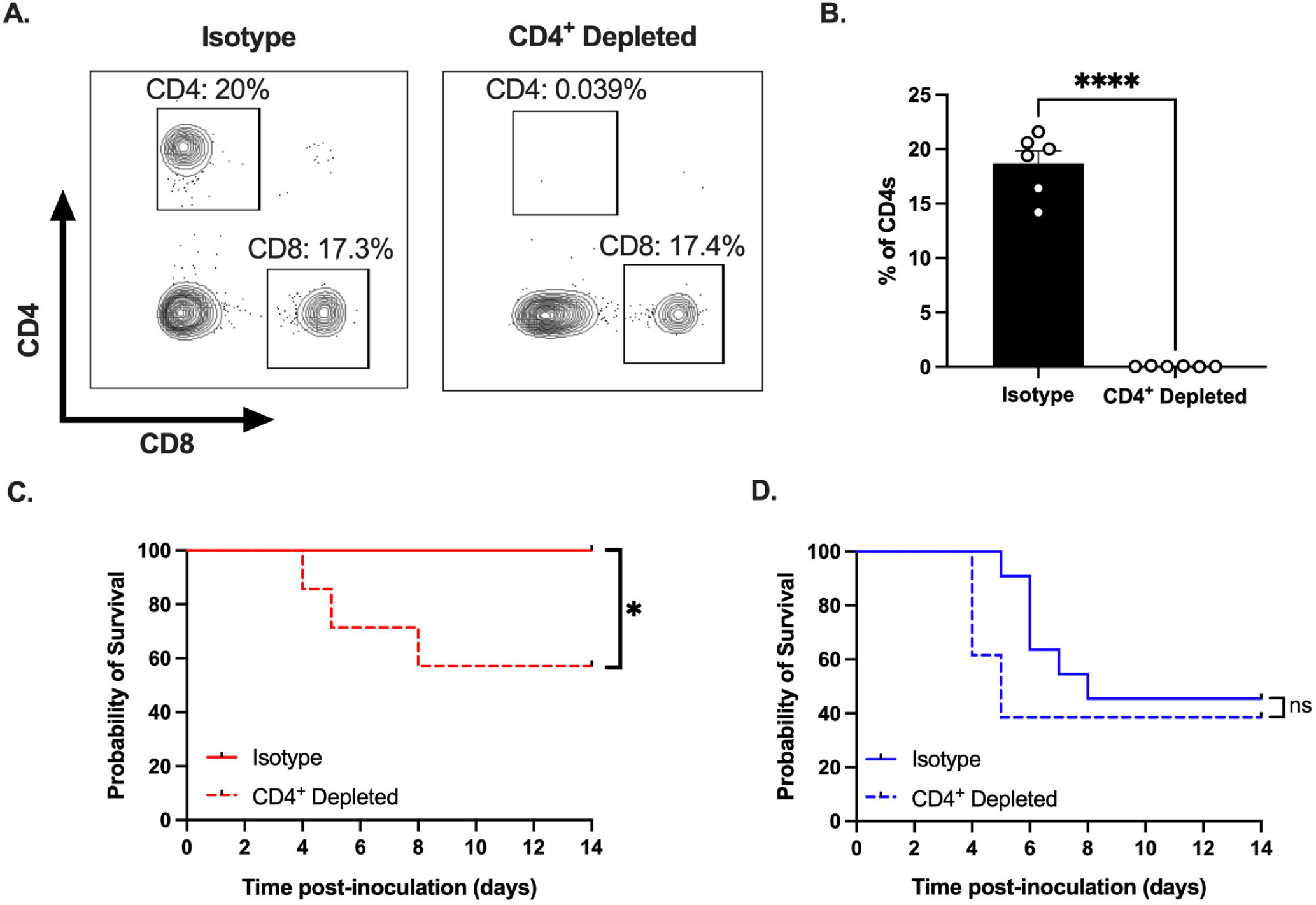
CD4^+^ T cells protect females from CVB3-induced lethality. (A) Representative flow cytometry plots of T cells gated on CD4^+^ and CD8^+^ T cell expression in female *Ifnar^-/-^*mice treated with an anti-CD4 antibody or isotype control. (B) The representative frequency of CD4^+^ T cells in female *Ifnar^-/-^* mice treated with an anti-CD4 antibody or isotype control from two independent experiments. ****p<0.0001, unpaired t-test. (C) Survival of CVB3-infected female mice treated with anti-CD4 or isotype control antibody. (D) Survival of CVB3-infected male mice treated with anti-CD4 or isotype control antibody. Survival data are collected from three independent experiments. n=11-14 mice per group. *p<0.05, Log-rank (Mantel-Cox) test.

## 3. Discussion

Previous research from our laboratory and others has demonstrated a strong sex bias in CVB3 pathogenesis (27, 30). The mechanisms dictating this sex bias are incompletely understood. Here, we found significantly more activated, viral-specific CD4^+^ T cells in infected female mice compared to male mice in the mesenteric lymph node and spleen as early as 5dpi. Moreover, the depletion of these CD4^+^ T cells protected female mice. These data provide evidence that activation and expansion of CD4^+^ T cells play a crucial role in limiting CVB3 pathogenesis and further demonstrate the importance that sex plays in immune protection against pathogens. The molecular mechanism for how CD4^+^ T cells protect against CVB3 is unclear, but sex contributes to CD4^+^ T cell effector function during different diseases and viral infections (18, 20, 21, 49-52). CD4^+^ T cells differentiate into multiple effector functions, including Th1, Th2, Treg, and Tfh cell types, which impact viral infections. We hypothesize that increased activation of CD4^+^ T cells promotes Th1 polarization in female mice, leading to cell-mediated viral clearance. Our previous studies and others on the role of sex hormones on Th1 differentiation may support this hypothesis. First, we also recently found that CD8^+^ T cells protect against CVB3 infection in females, suggesting a strong Th1 response (30). Second, estrogen can help upregulate the Th1 transcription factor, T-box protein expressed in T cells (T-bet), in murine splenocytes (53). Finally, estradiol-treated mice have higher levels of IFN-γ producing CD4^+^ T cells, indicating the development of a Th1 response (54). Interestingly, *in vivo* data for CVB3 infections differs. Removing endogenous testosterone by gonadectomy promotes a Th2 response and increases Tregs following CVB3 infection (21). Furthermore, unlike many autoimmune diseases, autoimmune-induced myocarditis following CVB3 infection reflects a male bias driven by a Th1 response. Contrarily, Th2s in female mice reduce mortality (18, 24, 55). Therefore, while we have not fully characterized the CD4^+^ T cell subtypes following infection, it is possible that CVB3 also drives a predominantly Th2 response, limiting inflammation in our model. Thus, based on this dichotomy, it will be essential to characterize the CD4^+^ T cell effector subpopulations and their kinetics following infection of CVB3 in our mouse model.

Previous studies using C57BL/6 mice to evaluate the T cell response revealed a limited expansion of CD4^+^ and CD8^+^ T cells (19, 45, 46). However, these studies exclusively utilized male mice. Our data corroborate these findings but show a significant increase in CD4^+^ T cells in infected females. Further, our data demonstrating that CD4^+^ T cells are protective in female mice differ from previous models. Previous studies show that CD4^+^ T cells can be detrimental, rather than protective, to disease. For example, adult male CD4^+^ knockout mice are less susceptible to CVB3-induced disease (19). However, the discrepancy in our data may be due to strain-specific immune differences. Here, we used C57BL/6 mice that produce acute viral myocarditis, while Balb/c models develop autoimmune-mediated myocarditis following CVB3 infection. The contrasting results may be due to different predominant Th subtypes in Balb/c mice and C57BL/6 mice. Since the balance between Th1 and Th2 responses plays a role in exacerbating or limiting inflammation in the heart, this difference may be an underlying factor for our contrasting results.

Another possible explanation for the differences between our data and other labs is our model’s absence of type I IFN. The interaction between CD4^+^ T cells and type I IFN is complex. On the one hand, type I IFN can promote Th1 differentiation (56, 57). Conversely, they may also be required for the Tfh differentiation (58). Moreover, in the context of the same infection, the kinetics of type I IFN signaling can have opposing effects on CD4^+^ T cell fates (59). The lack of type I IFNs could alter CD4^+^ T cell fates in male and female mice. Therefore, future work is required to understand how type I IFNs impact T cells in the context of CVB3 infections.

Finally, the mechanism for how CD4^+^ T cells protect female mice is under investigation. We hypothesize that a classical model of Th1 T cells likely primes the CD8^+^ T cell response for cell-mediated killing of virally infected cells. However, the CD8^+^ T cell response to CVB3 is limited in our model. Therefore, it is intriguing to speculate that cytotoxic CD4^+^ T cells may also play a role in limiting virally infected cells. These cytotoxic CD4^+^ T cells have been shown to limit viral infections in an MHC II-dependent mechanism (47). In addition to cytotoxic CD4^+^ T cells, newly recruited CD4^+^ intraepithelial lymphocytes in the intestine have also recently been shown to limit gut infection of adenovirus (60). The impact of cytotoxic CD4^+^ T cells and intraepithelial lymphocytes on CVB3 is unclear, but future studies may define their role in limiting CVB3 pathogenesis.

In summary, our data indicates that CD4^+^ T cells play a protective role in CVB3-induced mortality and protect female mice. These data highlight the importance of sex-dependent T cell responses to pathogens. While the mechanism is still unclear on what drives this sex bias in T cell response, ultimately uncovering these mechanisms will have essential implications in vaccine development to limit CVB3 infections.

## Data Availability statement

The original contributions presented in this study are included in the article or supplementary material. Further inquiries can be directed to the corresponding author.

## Ethics Statement

All animals were handled according to the Guide for the Care of Laboratory Animals of the National Institutes of Health. All mouse studies were performed at Indiana University School of Medicine (IUSM) using protocols (Approved IUSM Protocol: 20075) approved by the local Institutional Animal Care and Use Committee in a manner designated to minimalize pain, and any animals that exhibited severe disease were euthanized immediately by CO_2_ inhalation.

## Author Contributions

The study was designed by AP, AD, LS, SC, MR, and CR. AP, AD, SC, and CC performed the experiments. The results were interpreted by AP, AD, LS, SC, MR, and CR. AP and CR wrote the manuscript; AP, AD, LS, SC, and MR contributed to the manuscript preparation. All authors contributed to the article and approved the submitted version.

## Acknowledgments

We want to thank the members of the Indiana University Melvin and Bren Simon Cancer Center Flow Cytometry Resource Facility for their outstanding technical support. We thank the NIH Tetramer Core Facility (contract number 75N93020D00005) for providing the GP66 tetramer.

